# Proteomic signatures of sublethal deltamethrin exposure in the European earwig

**DOI:** 10.64898/2026.05.05.722842

**Authors:** Laura Pasquier, Daniel Tomas, Valérie Labas, Ana Paula Teixeira-Gomes, Joël Meunier, Charlotte Lécureuil

## Abstract

Pesticides are ubiquitous in agroecosystems and pose substantial risks to non-target organisms. Traditional ecotoxicological assessments focus on survival, reproduction, or overt behaviour, yet these endpoints may fail to detect subtle, molecular-level stress. Here, we investigated the effects of sublethal deltamethrin exposure on the head proteome of field-collected European earwig (*Forficula auricularia*) females, sampled at two life stages (pre-oviposition and post-family life) to account for physiological context. Our results reveal that deltamethrin induces a robust proteomic response shared across developmental stages, including the regulation of key detoxification enzymes (NADPH–cytochrome P450 reductase, arginine kinase). In parallel, stage-specific responses were observed, involving proteins related to metabolism, stress response, and cellular organization. Strikingly, these molecular perturbations occurred without detectable changes in reproductive traits, highlighting a disconnect between cellular stress and organismal phenotypes. Several uncharacterized proteins were consistently regulated, representing promising targets for future studies on pesticide adaptation and potential detoxification pathways. Overall, these findings suggest that classical phenotypic assays may underestimate sublethal pesticide effects, and that proteomic profiling provides a sensitive framework to uncover underlying molecular responses. By integrating natural variability, realistic exposure, and reproductive physiology, our study emphasizes the need for molecular approaches in environmental risk assessment and offers a new perspective on the subtle, cryptic effects of agrochemicals.

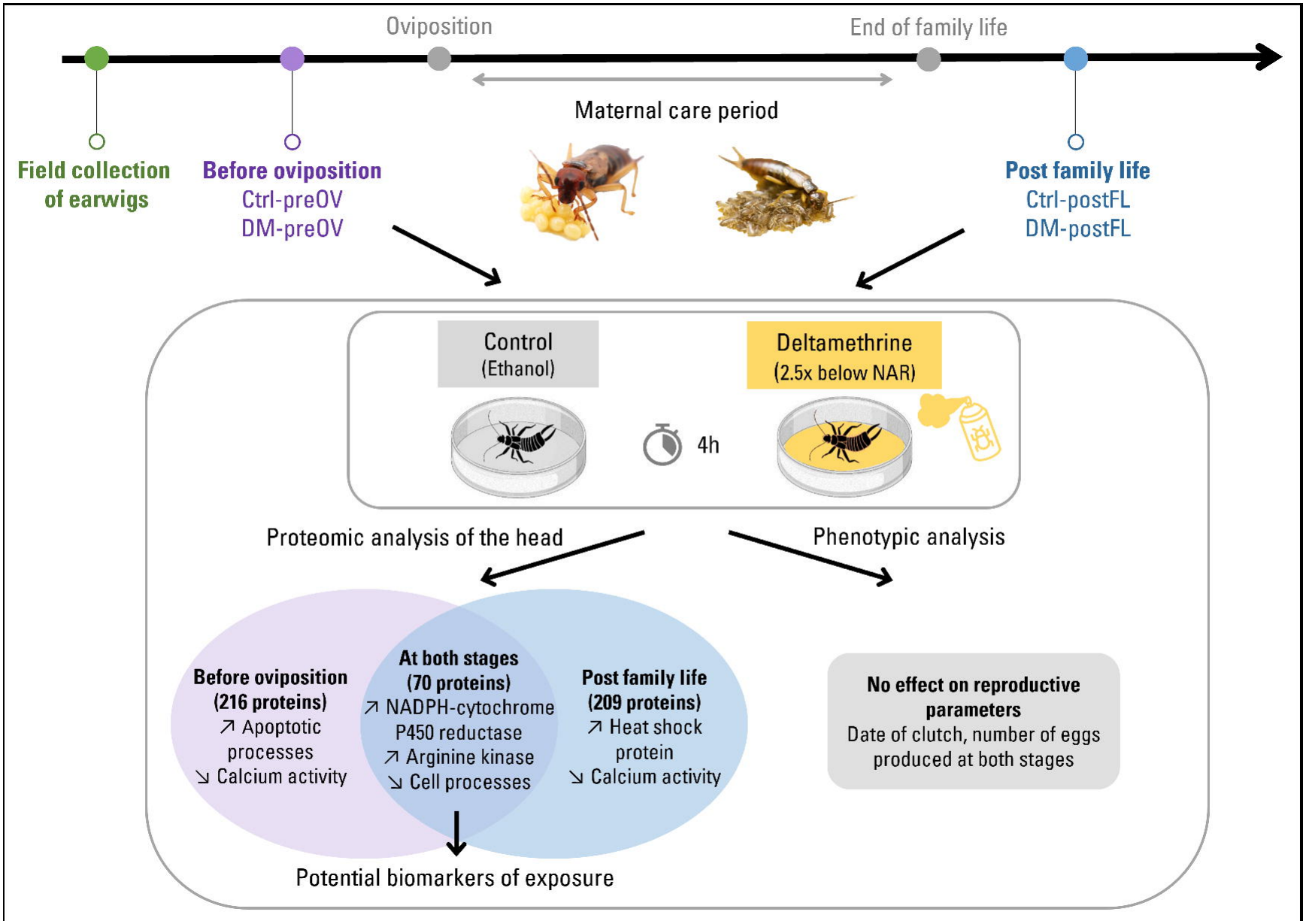

## 1. INTRODUCTION

Pesticides are among the most pervasive pollutants in modern agroecosystems, and their intensive use has become a major driver of environmental contamination worldwide. Although designed to control agricultural pests, pesticides inevitably affect a wide range of non-target terrestrial insects, many of which play essential ecological roles (Desneux et al. 2007; Emmerson et al. 2016). Ecotoxicological research has documented effects across a broad range of phenotypic traits, including survival, reproduction, but also sublethal phenotypic effects on behaviour and physiology, with potential consequences for individual fitness, population dynamics, and ecosystem functioning (Müller 2018; Feng et al. 2023). However, conventional phenotypic endpoints provide only a partial view of how pesticide exposure affects organisms. They provide limited insight into the biological mechanisms underlying observed effects and may constrain our ability to establish causal pathways and predict responses across species (Lemos et al. 2010; Canzler et al. 2020). Moreover, compensatory processes may maintain apparent phenotypic stability, despite underlying molecular or physiological perturbations (Diz et al. 2012; Martelli et al. 2020; Lesseur et al. 2023), while delayed, cumulative, or transgenerational effects may remain undetected when only a limited set of phenotypic traits is measured (Karlsson et al. 2021; Stuligross and Williams 2021).

To complement phenotypic endpoints, sublethal biomarkers have long been used in ecotoxicology to detect physiological and biochemical responses to chemical stress, including changes in specific processes such as detoxification, oxidative stress or neurotoxicity (Lagadic et al. 1994; Van Gestel and Van Brummelen 1996; Beliaeff and Burgeot 2002). More recently, omics approaches, and proteomics in particular, have broadened this perspective by providing a more comprehensive characterization of molecular responses to pesticide exposure, allowing changes across multiple biological pathways to be identified (Lemos et al. 2010; Canzler et al. 2020). Proteomic analyses can reveal molecular adjustments to pesticide exposure associated with processes such as metabolic reprogramming, cellular repair, or detoxification, including subtle biochemical alterations that may precede phenotypic effects or occur without measurable phenotypic consequences, thereby helping to identify candidate proteins and pathways for subsequent targeted investigation (Diz et al. 2012; Martelli et al. 2020). This broader molecular response has been observed, for instance, in *Drosophila melanogaster*, where sublethal imidacloprid exposure altered proteins associated with oxidative stress and retinal function without immediate behavioural changes (Martelli et al. 2020).

Deltamethrin is a widely used agricultural insecticide belonging to the pyrethroid class (Dietz et al. 2009; Rehman et al. 2014). It acts primarily on voltage-gated sodium channels causing neuronal hyperexcitation and, at sufficiently high exposure levels, paralysis and death (Laufer et al. 1984; Narahashi et al. 1992). However, responses to deltamethrin can vary considerably among species and biological traits, meaning that effects observed in standard toxicity assays may not fully capture its consequences for non-target organisms (Cheng et al. 2015; Zhang et al. 2022; Varela et al. 2024). This is particularly relevant for the European earwig (*Forficula auricularia*), a widespread omnivorous insect commonly found in orchards and gardens that can be exposed to insecticides in agricultural environments and has been shown to be sensitive to deltamethrin (Meunier et al. 2020; Mauduit et al. 2021). Its ecological importance as a predator contributing to biological control service in pip-fruit orchards further makes *F. auricularia* a relevant non-target species for assessing insecticide effects (Dib et al. 2011; Orpet et al. 2019; Alins et al. 2023). Moreover, *F. auricularia* exhibits an extended period of maternal care, with females remaining with their eggs for approximately 50 days and subsequently with newly hatched juveniles (nymphs) for about two weeks before potentially producing a second clutch (Meunier et al. 2012; Pasquier et al. 2025a). This prolonged association between mothers and offspring provides a distinctive biological context in which pesticide exposure may affect not only individual physiology and behaviour, but also interactions within family groups.

Recent studies have investigated the effects of exposure to various pesticides on phenotypic and behavioural traits in *F. auricularia* (Malagnoux et al. 2014, 2015a; Le Navenant et al. 2019; Meunier et al. 2020; Mauduit et al. 2021; Merleau et al. 2022; Pasquier et al. 2024, 2025b). In particular, deltamethrin exposure produces trait-specific effects, with some maternal behaviours being reduced while others remain unaffected (Meunier et al. 2020; Mauduit et al. 2021). Physiological and reproductive responses are similarly variable, with effects reported on clutch production and hatching success, but not on detectable effects on egg development, offspring weight, female weight gain, or survival(Meunier et al. 2020; Mauduit et al. 2021). At the molecular level, exposure to various pesticides can induce changes in detoxification enzymes, including glutathione S-transferases, carboxylesterases, and acetylcholinesterase, reflecting an adaptive metabolic response to chemical stress (Malagnoux et al. 2014; Le Navenant et al. 2019). However, whether deltamethrin elicits comparable or distinct responses at the proteomic level in *F. auricularia*, and how these responses relate to the variable phenotypic effects observed, remains unknown. Investigating the earwig proteome could therefore reveal early or subtle molecular responses to deltamethrin that are not detectable through conventional phenotypic endpoints, while providing insight into the mechanisms underlying its effects.

In this study, we characterized changes in protein abundance induced by sublethal exposure to deltamethrin in *F. auricularia* females. We focused on the head proteome because deltamethrin primarily targets neuronal sodium channels (Laufer et al. 1984; Narahashi et al. 1992). A total of 163 females were exposed to deltamethrin at 2.5× below the standard application rate in French orchards, and sampled at two distinct reproductive stages: before oviposition and post family-life. These stages differ markedly in reproductive investment and physiological state, with post-family-life females having depleted energy reserves following prolonged fasting and maternal care at low winter temperatures (Honorio et al. 2025). Such physiological differences are likely to influence pesticide susceptibility, providing an ecologically relevant framework to assess stage-specific proteomic responses. Of the 163 females, 36 were used for differential bottom-up proteomic analysis of the heads using a label-free quantitative approach. The remaining 127 females were monitored for reproductive performance, including day of oviposition and egg number. Overall, this experimental design allowed us to directly test three ecotoxicologically relevant hypotheses. First, sublethal deltamethrin exposure induces general molecular stress responses, consistently affecting protein profiles across reproductive stages. Second, physiological context modulates these responses, producing stage-specific changes before oviposition versus after maternal care. Third, sublethal exposure triggers diverse molecular alterations detectable at the proteomic level, revealing subtle effects that are not necessarily apparent in conventional phenotypic measures. Altogether, this approach integrates realistic environmental exposure, natural inter-individual variability, and reproductive physiology to provide a robust and ecologically meaningful assessment of pesticide impacts at the proteomic level.

## 2. MATERIALS AND METHODS

### 2.1 Experimental design

The experiment involved a total of 163 earwig females randomly sampled from approximately 2,000 males and females collected in July 2022 from an organic peach orchard near Valence, France (Lat 44.978526, Long 4.926383). All females belonged to the species *Forficula auricularia* Linnaeus, 1758, also known as *Forficula auricularia* clade A (González-Miguéns et al. 2020). After collection, adults were randomly assigned to large plastic containers with a balanced sex-ratio and maintained under standard laboratory conditions (18–20 °C, 12:12 h light:dark cycle). This setup allowed uncontrolled mating and natural group living until oviposition (Sandrin et al. 2015). When the first field-sampled female oviposited (approximately four months after collection), all non-ovipositing females were individually isolated in Petri dishes (diameter 5 cm) and kept in complete darkness at 10 °C to simulate natural winter subterranean conditions (Honorio et al. 2025). Females typically oviposit within a few days to a few weeks after isolation. From this pool of 1,000 isolated females, 163 were randomly selected for the present experiment (the remaining were used in other experiments not presented here).

One week after isolation, 79 of the 163 females were randomly assigned to either deltamethrin (N = 43, DM-preOV) or to a control (N = 36, Ctrl-preOV) treatment. Exposure followed a standard protocol designed to mimic realistic exposure to pesticide residues (Malagnoux et al. 2015a; Meunier et al. 2020; Pasquier et al. 2024, 2025b). In brief, 88 µL of deltamethrin solution (SIGMA #45423; 3 µg/mL; final surface dose 6.875 ng/cm², which is 2.5× below the normal application rate in French orchards) or ethanol (control) was evenly applied to the ground of a new Petri dish (diameter 5 cm). Each female was then gently transferred to the treated Petri dish and allowed to walk freely for 4 hours in the dark at room temperature before being returned to its initial Petri dish (Meunier et al. 2020; Mauduit et al. 2021; Pasquier et al. 2024, 2025b). The treated petri dishes were renewed for each female. Seven days later, heads from 18 females (9 females per treatment) were dissected for proteomic analysis. The choice of nine independent biological replicates per treatment and cohort was intended to maximise biological replication within the practical and cost constraints of the proteomic workflow. The remaining 61 females were monitored twice per week to record reproductive parameters, including the date of oviposition and, three days post-oviposition, the number of eggs produced.

The remaining 84 females were maintained under standard conditions until the end of family life, defined as 14 days after egg hatching (Honorio et al. 2025). They were kept in complete darkness at 10 °C until egg hatching. On that day, they were transferred with their full clutch of nymphs to a new Petri dish, and then maintained at 18-20°C under a 12:12 h light:dark cycle. Fourteen days later, 44 randomly selected females were exposed to deltamethrin (DM-postFL) and 40 to control solution (Ctrl-postFL) using the same protocol as above. Seven days later, heads were dissected for proteomic analysis (9 females per treatment), while the remaining females were monitored daily to record reproductive parameters, including second clutch production, the date of oviposition, and the number of second clutch eggs produced.

Throughout the experiment, each Petri dish was lined with moist sand. Pre-ovipositing females, and mothers with juveniles were provided with a standard laboratory diet mainly composed of pollen, cat food and bird seed (details in Kramer et al. 2015), renewed weekly. No food was provided between oviposition and egg hatching, as females typically cease during this period (Kölliker 2007).

### 2.2 Preparation of protein extracts

#### 2.2.1 Protein extraction

Female heads were dissected and immediately processed for protein extraction in a 2 mL Eppendorf tube. Each sample containing one female head was lysed in 100 µL of lysis buffer containing 10 mM Tris-HCl, pH 7.4, 2% SDS, protease inhibitor cocktail (Sigma, ref. P2714), added immediately before use (1 µL per sample). Mechanical disruption was performed using a Tissue Lyser (Qiagen, Hilden, Germany) and 1 Tungsten 7mm beads (Qiagen, Hilden, Germany) for 1 min at maximum speed. Samples were centrifuged at 11,000 × g for 10 min at 4 °C. The supernatant containing soluble proteins was collected and stored at −20 °C until further analysis.

#### 2.2.2 Protein quantification (BCA Assay)

Protein concentration was determined using a BCA assay kit (Uptima, Interchim, ref. UP40840A) according to the manufacturer’s instructions. For each sample, 1 µL of supernatant was loaded in triplicate into a 96-well microplate (Greiner 96 Flat transparent) and quantified against a BSA standard curve (0–2 mg/mL) prepared in lysis buffer (25 µL per well). Absorbance was measured at 595nM after incubation with 200 µL of BCA working reagent in each well at 37°C for 30min, using a Tecan microplate reader (i-control I12 software) and samples exceeding the standard range were diluted 1:10 and reassayed. Protein amounts ranged from 14.1mg of protein/g of head and 31.5mg of protein/g of head.

#### 2.2.3 Quality control of protein extracts

Protein extract quality was assessed by SDS-PAGE (Sodium Dodecyl Sulfate Polyacrylamide Gel Electrophoresis). Twenty micrograms of protein were loaded onto 4–20% gradient mini-gels (Bio-Rad, ref. 4561094). Prior to loading, samples were sonicated briefly and mixed with an appropriate volume of Laemmli buffer containing 0.0625 M Tris-HCl, pH 6.8, 2% SDS, 10% glycerol in presence of 5% TCEP (tris(2- carboxyéthyl)phosphine) and bromophenol blue. Samples were heated at 100°C for 5 min and centrifuged for 1 min before loading. Samples and Bio-Rad protein markers Precision Plus Protein™ Dual Color Standards were fractionated at 200 V in 1X running buffer containing 25 mM Tris base, 192 mM Glycine and 0.1% SDS. Following separation, gels were stained overnight with Coomassie Brilliant Blue R-350 (1:10 dilution in staining solution composed of 600 mL deionized water, 300 mL ethanol, and 100 mL acetic acid), using 50 mL per gel under gentle agitation (10 rpm). Destaining was performed in the same solution (without dye) until a clear gel was obtained. Gels were rinsed with water and stored in 2% acetic acid. The quality of the gels was confirmed by the presence of distinct protein bands, indicating that the proteins were not degraded.

#### 2.2.4 Sample preparation for “One-Band” gel analysis

For each sample, 25 µg of protein were deposited on a 10% polyacrylamide gel. Sample preparation was performed as described above. The electrophoretic migration was limited and stopped as soon as the bromophenol blue front entered the resolving gel, allowing all proteins to concentrate in a single band. The entire protein-containing band was excised and subdivided into small pieces. Gel fragments were stored at −80 °C until in-gel digestion for proteomic experiments.

### 2.3 High-Resolution Mass Spectrometry analysis

#### 2.3.1 Sample preparation: in-gel digestion

Gel pieces were washed in water:acetonitrile solution (1:1) for 5min followed by 100% acetonitrile for 10min. Protein reduction and cysteine alkylation were performed by successive incubations with 10 mM dithiothreitol in 50 mM NH_4_HCO_3_ for 30min, at 56°C, then 55 mM iodoacetamide in 50 mM NH_4_HCO_3_ for 20 min, in dark at room temperature. Gel pieces were then incubated with 50 mM NH_4_HCO_3_ and acetonitrile (1:1), for 10 min followed by 100% acetonitrile for 15min. Proteolytic digestion was carried out, overnight using a solution of trypsin (Sequencing grade, Roche diagnostics, Paris, France) at 12.5ng/μl in 25 mM NH_4_HCO_3_. Firstly, peptides were extracted by sonication and incubation in 5% formic acid. The supernatant was removed and saved for each step. Gel pieces were incubated twice in water:acetonitrile (1:1) in presence of 1% formic acid, for 10 min and in acetonitrile, for 5 min. These three peptide extractions were pooled and dried using a SPD1010 speedvac system (Thermosavant, Thermofisher Scientific, Bremen, Germany).

#### 2.3.2 Evotip sample preparation

Dried peptides samples were resuspended in 500µL buffer A (100% water in presence of 0.1% formic acid, Sigma) and sonicated 10 min. To purify and concentrate peptides, the sample aliquots were manually loaded onto EvoTip Pure tips (Evosep EV2013, 10x96 tips) packed with ReproSil-Pur C18 beads. In brief, according to the manufacturer’s instructions, Evotips were washed with 20μL buffer B (100% acetonitrile in presence of 0.1% formic acid, Sigma), and centrifuged for 60 sec at 800 x g. Tips were soaked in 2-propanol (Thermo Fisher Chemical) for several min until all are pale white. For equilibration, 20 μL of buffer A was added onto the Evotips and centrifuged for 60 sec at 800 x g. Each tip was loaded with 100 μL of sample and centrifuged for 60 seconds at 800 x g, followed by a wash using 20μL of buffer A and a centrifugation for 60 sec at 800 x g. Finally, 100 μL of buffer A was transferred onto each Evotip, which was then centrifuged for 10 sec at 800 x g to ensure the C18 resin remained wet. The Evotip box was filled with buffer A up to 1/3 volume for storage, at room temperature, during the peptide separations by the Evosep One liquid chromatography system (Evosep, Odense, Denmark) controlled by Chronos Software (version 5.8.3).

#### 2.3.3 Evosep separation

Peptides were separated on a 15 cm C18 Endurance analytical LC column (1.9µm ReproSil Saphir C18 beads, ID 150µm x 15cm, EV-1106) utilizing the standardized 15 samples-per-day (SPD) method, which is a 88-minute active LC gradient with a total run time of 92 minutes. The analytical column was equilibrated, at room temperature at 1500 nL/min. The gradient was obtained by flow injection of Evosep solvent A (100% H2O acidified with 0.1% formic acid, Sigma) and solvent B (100% acetonitrile acidified with 0.1% formic acid, Sigma). Peptides were eluted, at a flow rate of 220 nL/min, by a linear gradient up to 35% solvent B. The column was connected to an integrated ZDV liquid junction adapter (EV-1085) with a 20 µm ID fused silica emitter (EV 1087). The Evosep column was mounted on a nano-spray Flex NG™ source (Thermo Fisher Scientific, Bremen, Germany) combined to ion mobility mass spectrometry (IMS) using FAIMS (high-Field Asymmetric waveform Ion Mobility Spectrometry).

#### 2.3.4 NanoLC-MS/MS

The Tribrid Orbitrap Ascend mass spectrometer (Thermo Fisher Scientific, Bremen, Germany) was operated in DDA (Data Dependent Acquisition) and in positive ion modes, automatically switching between MS and MS/MS acquisition for a 3 s cycle time. The source voltage was 1.9 kV, the capillary temperature was at 275°C and S-lens at 60%. Full-scan MS profile spectra (1 microscan) were acquired from 350 to 1400 m/z mass range in the Orbitrap, with a maximum injection time set to Auto with an absolute automatic gain control value set at 4.10^5^ ions. Resolution in the Orbitrap analyzer was set at R=240,000 (at m/z 200). Monoisotopic peak determination (MIPS) was performed in peptide mode on the most abundant peaks, presenting 2-7 charges and a minimum intensity threshold at 5.10^3^. The precursor ions were filtered using the quadrupole with an isolation window of 1.6 m/z (without isolation offset) and isolated in the linear ion trap. In the high-pressure cell of the ion trap, HCD (higher energy collisional dissociation) fragmentation mode was performed with a fixed normalized collision energy at 30%. MS2 centroid spectra (1 microscan) were acquired using the Rapid ion trap scan rate ranged from 200-1400 m/z using a maximum injection time in mode auto and a standard normalized AGC target for an absolute AGC value at 1.10^4^. Dynamic exclusion was activated during 60 seconds with a repeat count of 1, using a mass tolerance of 10 ppm (high and low), excluding also the corresponding isotopes. A lock mass correction was enabled for accurate mass measurements using the run-start Easy-IC™ mode using fluoranthene cations (at 202.07770 m/z).

For all nanoLC-FAIMS-MS/MS experiments, FAIMS Pro Duo interface (Thermo Fisher Scientific, San Jose, CA) was installed and set to standard resolution. The distance between the silica emitter and the FAIMS inlet was controlled to 2-3 mm. The nitrogen carrier gas was set as optimized 4.0 L/min.

For each sample, proteomic experiments were performed in duplicate using 2 different IMS methods with 3 constant compensation voltages (CVs). Ion fractionation was applied using the combination -35, -50, -70 CVs, for the method or -45, -60, -75 CVs, allowing transmission of specific groups of ions for 1.2 sec, 1 sec and 0.8 sec, respectively in that order.

### 2.4 Statistical analysis

Acquired Orbitrap Ascend. raw files were processed with Proteome Discoverer 3.1 software (ThermoFisher Scientific, Bremen, Germany). Samples designated as Ctrl-preOV and DM-preOV in the manuscript corresponded to Ctrl1 and DM1 in the raw data files, respectively, while Ctrl-postFL and DM- postFL corresponded to Ctrl2 and DM2. MS/MS ion searches were performed using the SEQUEST HT search engine against a UniprotKB database Insecta (Taxonomy_id=50557 version 2024-1002). The search parameters included trypsin as a protease with two allowed missed cleavages. Carbamidomethylcysteine was set as static modifications, and methionine oxidation, loss of methionine and/or acetylation of N-term protein as variable modifications. Ion tolerances were 5 ppm for parent ions and 0.6 Da for fragment ions. SEQUEST results from target and decoy database searches were filtered using a False Discovery Rate (FDR) of 0.01 for proteins and peptides (minimum 6 amino acids). For comparative analyses, protein abundance ratios between groups (pairwise ratio-based method) were calculated after normalization to total peptide abundance, excluding modified peptides. A background-based t-test was applied to assess significance, comparing individual peptide/protein ratios against the global background distribution. P-values were adjusted using Benjamini–Hochberg correction. Subsequent statistical analyses were performed in R v4.1.1 (https://www.r-project.org/) using the packages *topGO* (Alexa and Rahnenführer 2006). Differentially expressed proteins were first identified across all developmental stages with log2 abundance ratio ≥ 1 and a P-value ≤ 0.05 for upregulated proteins, or log2 abundance ratio ≤ –1 and a P-value ≤ 0.05 for downregulated proteins. Gene Ontology (GO) enrichment analyses were then conducted separately for up-and downregulated proteins using the *topGO* package. For each comparison, a binary protein list indicating the presence of differentially expressed proteins among all identified proteins was constructed, and proteins mapped to their associated GO terms. Enrichment was tested within the Biological Process ontology using Fisher’s exact test combined with the *weight01* algorithm, which accounts for the hierarchical structure of GO terms. Significantly enriched terms (p ≤ 0.05) were extracted and ranked by Fisher’s statistic. In addition, stage-specific analyses were conducted for Ctrl-preOV vs DM-preOV and Ctrl-postFL vs DM-postFL. For each stage, up- and downregulated proteins were identified. GO terms associated with these stage-specific proteins were extracted and analysed using the same *topGO* approach, providing insight into molecular pathways consistently or differentially affected at each developmental stage. For the phenotypic parameters, we analysed the time to oviposition using a Cox proportional hazards regression model, which accounts for censored data from females that did not produce a clutch. The number of eggs produced in the first and second clutches was analysed using two linear models (*lm* function R).

## 3. Results

In total, 1,532 proteins were identified (with FDR < 1%) in the heads of *F. auricularia* females across all samples and reproductive stages, including 128 without GO terms. Overall, deltamethrin exposure altered distinct sets of proteins, with some showing consistent responses across both pre-oviposition and post-family life stages, and others showing stage-specific responses.

The consistent proteomic response comprised 70 proteins (Log2 Fold Change ≥ 1 or ≤ -1), of which 44 were upregulated and 26 downregulated (Figure 1.A & 1.B; Table III). Upregulated proteins included NADPH–cytochrome P450 reductase (detoxification), arginine kinase (energy metabolism), and glutamate dehydrogenase (neuroprotection), while downregulated proteins included short-chain specific acyl-CoA dehydrogenase (lipid metabolism) and the heat-shock proteins HSP70 and HSP83 (cellular stress responses). GO enrichment of the 41 upregulated proteins (6.8% lacked GO annotations) revealed four overrepresented biological processes: acylglycerol metabolic process, regulation of canonical Wnt signaling pathway, negative regulation of Wnt signaling pathway, and cell cycle. This indicates enhanced energy buffering, cellular signaling and proliferation (Table II). Conversely, enrichment analysis of the 25 downregulated proteins (3.8% lacked GO annotations) identified four overrepresented processes: cytoskeleton-dependent cytokinesis, mitotic cell cycle process, cell cycle and cytoskeleton organization. This suggests reduced proliferative capacity and impaired cytoskeletal dynamics (Table II).

**Figure 1.**
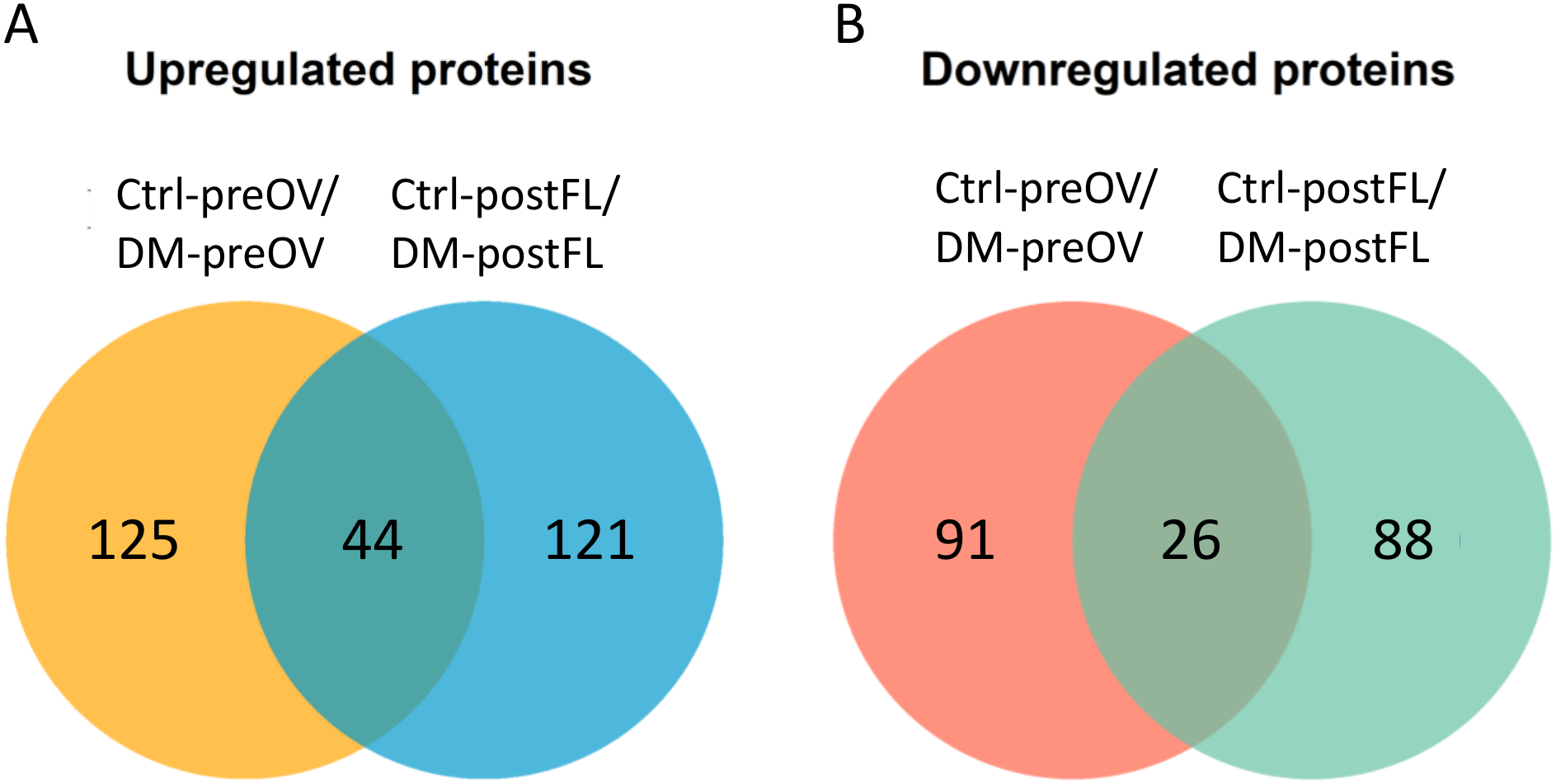
Number of (A) upregulated and (B) downregulated proteins in the head of females following deltamethrin exposure at both pre-oviposition (Ctrl-preOV/DM-preOV) and post-family life (Ctrl-postFL/DM-postFL) stages.

**Table II.**
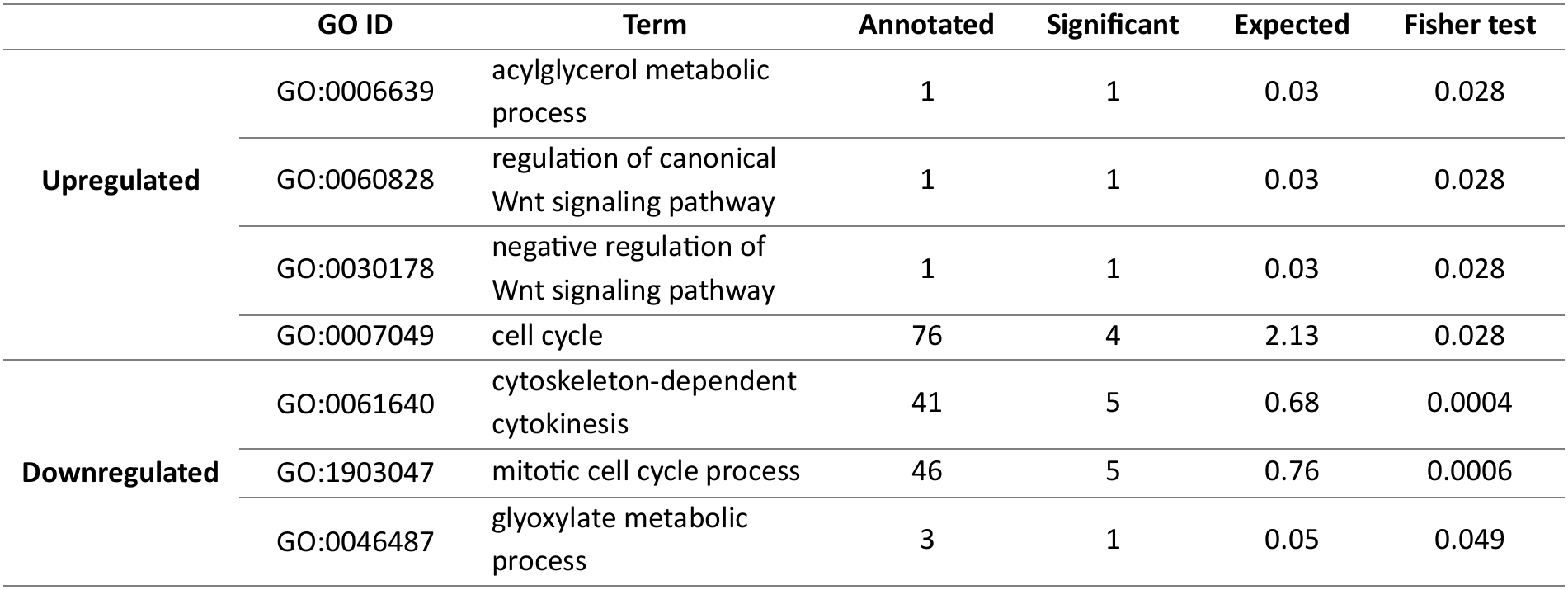
GO term enrichment specific to deltamethrin treatments on the 70 proteins differentially expressed across both pre-oviposition and post-family life stages. GO term enrichment was considered significant at p ≤ 0.05, indicating biological process categories that are overrepresented among the upregulated or downregulated proteins compared to the full set of 1,532 identified proteins.

**Table III.** List of the 70 proteins specific to deltamethrin treatments across both pre-oviposition and post-family life stages. Upregulated proteins are defined by a log₂ fold change ≥ 1 and an adjusted p-value ≤ 0.05 and downregulated proteins by a log₂ fold change ≤ –1 and an adjusted p-value ≤ 0.05. The accession numbers were obtained from UniprotKB database from Insecta taxonomy.

|  | N° accession | Description |
|---|---|---|
| **Downregulated** | A0A5N4AMS7 | Actin, cytoplasmic |
|  | A0A2J7PWC8 | Tubulin alpha chain |
|  | J7KMD7 | Actin (Fragment) |
|  | Q8T0X3 | Putative actin (Fragment) |
|  | A0A2Z5E4L2 | Beta-actin (Fragment) |
|  | A0A9N9S3H6 | Actin |
|  | A0A2S2PCQ5 | Actin, muscle (Fragment) |
|  | T1HWW5 | isocitrate dehydrogenase (NADP(+)) (Fragment) |
|  | A0A9R1TUV1 | Talin-2 isoform X1 |
|  | Q6QIY6 | Sex-lethal |
|  | Q9U8K2 | Peptidyl-prolyl cis-trans isomerase (Fragment) |
|  | A0A3L8DUZ4 | Short-chain specific acyl-CoA dehydrogenase, mitochondrial |
|  | A0A8J6LAL0 | Twitchin |
|  | A0A2A4JVT8 | Twitchin |
|  | A0A6J0RP38 | Twitchin |
|  | A0A2H8TN08 | cAMP-dependent protein kinase catalytic subunit |
|  | E9JS33 | Sodium/potassium-transporting ATPase subunit alpha (Fragment) |
|  | A0A2H1VPY4 | SFRICE_000162 |
|  | A0A3Q0J343 | Alpha-actinin, sarcomeric |
|  | A0AA96S1S4 | Heat shock protein 70 (Fragment) |
|  | A0A9P0H3J9 | ATP synthase subunit alpha |
|  | A0A1Z1JMZ2 | Heat shock protein 83 |
|  | A0A9P0G9Y4 | vesicle-fusing ATPase |
|  | A0A1B0EXR1 | LIM zinc-binding domain-containing protein |
|  | A0A1L8DMV3 | Putative regulatory protein mlp |
|  | A0A9N9RMV4 | 2-oxoglutarate dehydrogenase, mitochondrial |
| **Upregulated** | A0A194Q983 | Uncharacterized protein |
|  | A0A9P0H4Y6 | Titin |
|  | A0AAG5DRS1 | Uncharacterized protein |
|  | A0A6A4ISF6 | 60S acidic ribosomal protein P0 |
|  | A0A8B8I4T8 | Myosin heavy chain, muscle isoform X44 |
|  | A0A146MEA6 | Myosin heavy chain, non-muscle (Fragment) |
|  | A0A6P7GE44 | V-type proton ATPase subunit H isoform X1 |
|  | A0A8B8HJ34 | Tubulin beta chain |
|  | A0A6M2DDS7 | Tubulin alpha chain (Fragment) |
|  | A0A9E9EP55 | Tubulin alpha chain |
|  | A0A8D8MX96 | Actin, muscle (Fragment) |
|  | A0AAJ7V0F3 | 26S proteasome regulatory subunit 8 |
|  | A0A2J7RK34 | Talin-1 |
|  | A0A3Q0J2Q1 | Uncharacterized protein LOC103513782 |
|  | A0A336KP24 | Elongation factor 1-alpha |
|  | A0A6L2Q096 | Ras-related protein Ral-a |
|  | A0A0M3R7Y6 | NADPH--cytochrome P450 reductase |
|  | A0A6J2YV18 | Acetyl-CoA carboxylase isoform X1 |
|  | A0A1B1VWB3 | arginine kinase (Fragment) |
|  | A0A0C5DQD1 | arginine kinase (Fragment) |
|  | A0A2J7PZH5 | Diacylglycerol kinase |
|  | A0A7R9IPI7 | Dihydrolipoyl dehydrogenase |
|  | A0AAD7YCY8 | Glucose-6-phosphate isomerase |
|  | A0A8I6RK91 | glutamate dehydrogenase [NAD(P)(+)] |
|  | A0A1L6Z4U2 | Isocitrate dehydrogenase (Fragment) |
|  | A0A7R9D8K9 | Protein kinase domain-containing protein |
|  | A0A482XTB1 | Protein kinase domain-containing protein |
|  | A0A0P4VQK4 | Pyruvate dehydrogenase E1 component subunit alpha (Fragment) |
|  | A0A182F344 | Rab GDP dissociation inhibitor |
|  | A0A8S1CPN3 | Integrin alpha-2 domain-containing protein |
|  | A0A0A9X8I3 | 26S proteasome non-ATPase regulatory subunit 11 (Fragment) |
|  | A0A2L0CAN6 | 60A calcium-transporting ATPase (Fragment) |
|  | A0A2L0CA80 | 60A calcium-transporting ATPase (Fragment) |
|  | A0A8D9ALE4 | Dystonin |
|  | A0A6M2DCX7 | Putative ca2+-binding actin-bundling protein |
|  | A0A7R9CKQ9 | ATP synthase subunit alpha |
|  | B4JUR1 | GH15224 |
|  | Q9NGH2 | Heat shock cognate protein 70 (Fragment) |
|  | A0A151I798 | Protein NipSnap |
|  | A0A977MTF8 | Prohibitin (Fragment) |
|  | A0AAD4K2S2 | NADH dehydrogenase [ubiquinone] flavoprotein 1, mitochondrial (Fragment) |
|  | A0A195CU61 | 26S proteasome non-ATPase regulatory subunit 7 |
|  | A0A8D8HNY4 | Protein mago nashi |
|  | A0A1Y1LI32 | V-type proton ATPase subunit |

Proteins whose abundance differed significantly between deltamethrin-exposed and control females at only one life stage were identified, comprising 216 proteins in pre-oviposition females and 209 in post-family life females. In the pre-oviposition females, 125 of the 216 proteins were upregulated and 91 downregulated. Of the upregulated proteins (DM-preOV vs. Ctrl-preOV, Figure 1.A) (Log2 FC ≥ 1 or ≤ –1), 117 (94%) were successfully annotated and included in a GO enrichment analysis, which revealed 11 significantly enriched biological processes (Table IV). These processes were predominantly related to fatty acid metabolism, cellular organization, signal transduction, with a marked upregulation of apoptotic processes. Notably, 100 of these 117 proteins were exclusively detected after exposure, being completely absent beforehand. These newly appearing proteins were predominantly associated with metabolic and cellular processes (e.g., 2-oxoglutarate dehydrogenase, 26S proteasome non-ATPase subunit, clathrin heavy chain, medium-chain specific acyl-CoA dehydrogenase), intracellular transport (calcium-transporting ATPase), and muscle function (myosin tail domain-containing protein, paramyosin, tubulin alpha/beta chain) (Supplementary Data – Table S1). Of the 91 downregulated proteins detected before oviposition (DM-preOV vs. Ctrl-preOV, Figure 1.B) (Log2 FC ≥ 1 or ≤ –1), 77 (88%) were successfully annotated and included in a GO enrichment analysis, which revealed nine significantly enriched biological processes (Table IV). These processes were mainly associated with muscle structure, including myofibril assembly, calcium ion homeostasis, and calcium transport and lipid process. Interestingly, among these downregulated proteins, 63 were present in controls but absent in deltamethrin-exposed females, most of which were linked to muscle function (e.g., tubulin alpha chain, muscle LIM protein), calcium regulation (calcium-transporting ATPase), and cellular activity.

**Table IV.** GO term enrichment among the 216 differentially expressed proteins between Ctrl-preOV and DM-preOV compared to all proteins at the pre-oviposition stage. GO term enrichment was considered significant at p ≤ 0.05, indicating biological process categories that are overrepresented among the upregulated or downregulated proteins compared to the full set of 1,532 identified proteins.

|  | GO ID | Term | Annotated | Significant | Expected | Fisher test |
|---|---|---|---|---|---|---|
| **Upregulated** | GO:0032271 | regulation of protein polymerization | 25 | 4 | 2.17 | 0.0007 |
|  | GO:0051791 | medium-chain fatty acid metabolic process | 4 | 3 | 0.35 | 0.0024 |
|  | GO:0006418 | tRNA aminoacylation for protein translation | 2 | 2 | 0.17 | 0.0074 |
|  | GO:0008064 | regulation of actin polymerization or depolymerization | 24 | 3 | 2.08 | 0.0076 |
|  | GO:0006810 | transport | 288 | 33 | 24.95 | 0.012 |
|  | GO:0030029 | actin filament-based process | 156 | 19 | 13.51 | 0.013 |
|  | GO:0044085 | cellular component biogenesis | 174 | 19 | 15.07 | 0.0165 |
|  | GO:0007010 | cytoskeleton organization | 211 | 25 | 18.28 | 0.0167 |
|  | GO:2001233 | regulation of apoptotic signaling pathway | 12 | 3 | 1.04 | 0.021 |
|  | GO:0009967 | positive regulation of signal transduction | 3 | 2 | 0.26 | 0.021 |
|  | GO:0043065 | positive regulation of apoptotic process | 3 | 2 | 0.26 | 0.021 |
| **Downregulated** | GO:0070588 | calcium ion transmembrane transport | 41 | 9 | 2.27 | 0.0008 |
|  | GO:0006874 | intracellular calcium ion homeostasis | 5 | 3 | 0.28 | 0.0085 |
|  | GO:0035088 | establishment or maintenance of apical/basal cell polarity | 3 | 2 | 0.17 | 0.0087 |
|  | GO:0006629 | lipid metabolic process | 37 | 3 | 2.05 | 0.017 |
|  | GO:0030003 | intracellular monoatomic cation homeostasis | 59 | 10 | 3.26 | 0.021 |
|  | GO:0055074 | calcium ion homeostasis | 37 | 8 | 2.05 | 0.024 |
|  | GO:0007049 | cell cycle | 76 | 8 | 4.2 | 0.037 |
|  | GO:0030239 | myofibril assembly | 46 | 6 | 2.54 | 0.037 |
|  | GO:0007165 | signal transduction | 86 | 7 | 4.75 | 0.045 |

In the post-family life females, deltamethrin exposure affected 209 proteins (fold change ≥ ±2; log₂FC ≥ 1 or ≤ –1), of which 121 were upregulated and 88 downregulated (DM-postFL vs. Ctrl-postFL, Figure 1.A, 1.B). GO term enrichment analysis conducted on the 108 annotated upregulated proteins (10.7% lacked GO annotation) revealed a single significantly overrepresented biological process: protein folding (Table V). Among these upregulated proteins, 93 were uniquely detected in deltamethrin-exposed females and were mainly associated with oxidative stress (heat shock protein cognate 5, heat shock cognate 71 kDa protein, heat shock protein 105 kDa), cell adhesion (coronin, septin), muscle function (myosin motor/tail domain/heavy chain, tubulin alpha/beta chain), and cytoskeletal organization (actin, actin-related protein 3). Of the 88 downregulated proteins at this stage, 79 (89.8%) were successfully annotated, revealing four significantly enriched biological processes related to proteolysis, calcium homeostasis and transport, and monatomic cation homeostasis (Table V). Notably, a 62 (71%) of these proteins were present in controls samples but absent in deltamethrin-treated females. These proteins were primarily involved in reproductive processes (insulin-like growth factor 2 mRNA-binding protein 1), calcium signaling and muscle activity (calcium-transporting ATPase, actin-related protein 3, tubulin alpha chain), and metabolic functions (phosphoglycerate kinase, medium-chain specific acyl-CoA dehydrogenase, 26S proteasome non-ATPase) (Supplementary Data – Table S2).

**Table V.** GO term enrichment among the 209 differentially expressed proteins between Ctrl-postFL and DM-postFL at the post-family life stage. GO term enrichment was considered significant at p ≤ 0.05, indicating biological process categories that are overrepresented among the upregulated or downregulated proteins compared to the full set of 1,532 identified proteins.

|  | GO ID | Term | Annotated | Significant | Expected | Fisher test |
|---|---|---|---|---|---|---|
| **Upregulated** | GO:0006457 | protein folding | 76 | 11 | 5.64 | 0.021 |
| **Downregulated** | GO:0030003 | intracellular monoatomic cation homeostasis | 59 | 9 | 3.41 | 0.013 |
|  | GO:0051603 | proteolysis involved in protein catabolic process | 28 | 4 | 1.62 | 0.018 |
|  | GO:0070588 | calcium ion transmembrane transport | 41 | 7 | 2.37 | 0.02 |
|  | GO:0055074 | calcium ion homeostasis | 37 | 6 | 2.14 | 0.032 |

In contrast to proteins showing consistent responses across stages or stage-specific effects, a subset of proteins exhibited stage-dependent but opposing responses to deltamethrin exposure. Among the proteins whose abundance was significantly affected by deltamethrin at both stages, 46 exhibited opposite patterns of regulation between stages. Of these, 27 were upregulated before oviposition but downregulated in the post-family life, whereas 19 showed the reverse pattern. Analysis of this subset did not reveal additional enriched biological processes beyond those identified in the stage-specific analyses.

Finally, although deltamethrin exposure affected proteomic profiles, it did not influence reproductive timing or output. In females exposed before oviposition, no effects were observed on the laying date (Figure 2.A; F_1,59_= 0.91, P = 0.343) or the number of eggs produced (Figure 2.B; F_1,58_= 0.90, P = 0.347). Moreover, no second clutch was produced by females exposed at the end of family life, regardless of deltamethrin exposure (Figure 2.C) and number of eggs produced in the second clutch (Figure 2.D).

**Figure 2.**
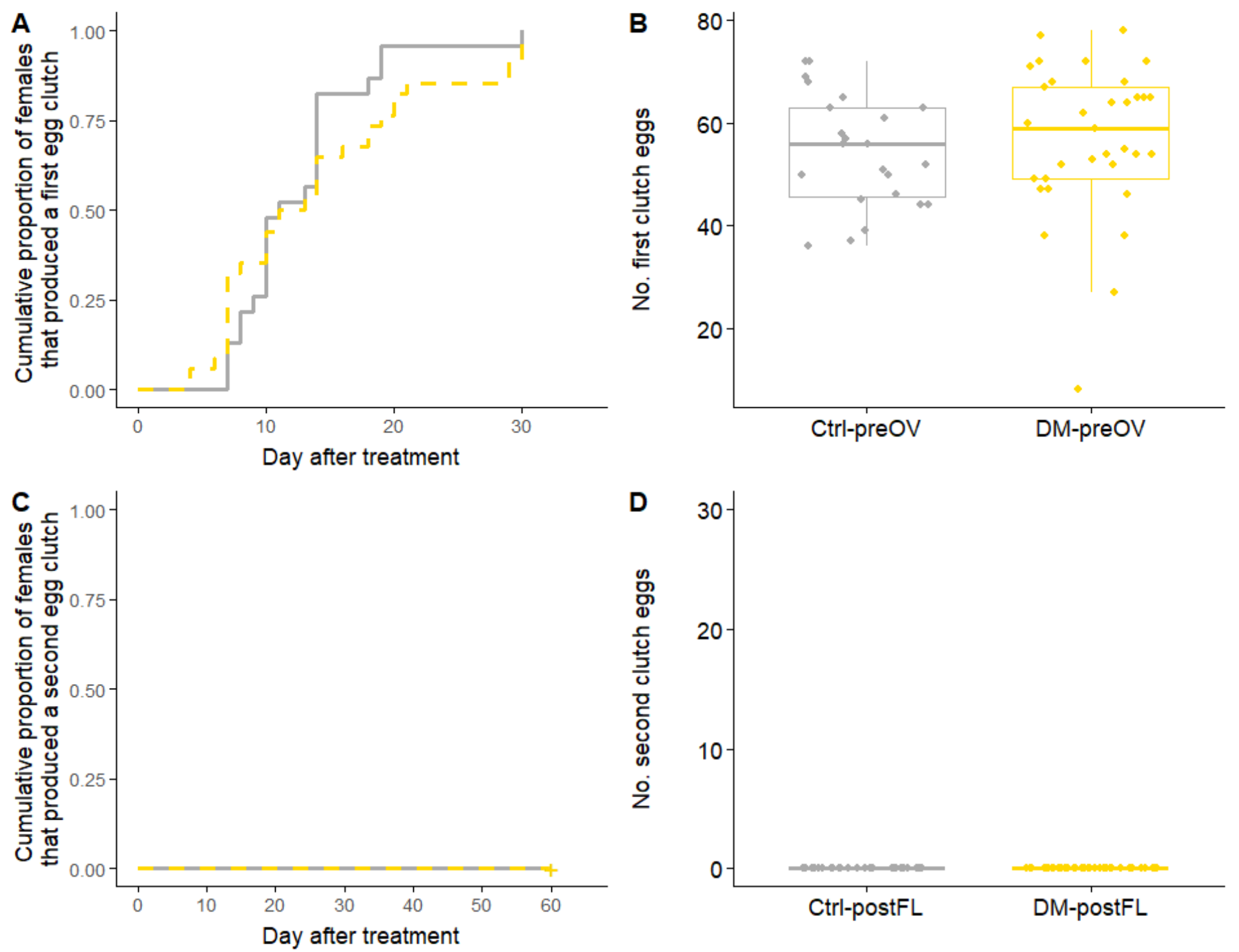
Effect of deltamethrin on reproductive parameters. A) Cumulative proportion of females that produced a first clutch of eggs starting four days after treatment, and B) number of eggs produced by females in the pre-oviposition treated females (Ctrl-preOV, N=27 and DM-preOV, N=34; all females are shown, as all females included in the analysis laid eggs). C) Cumulative proportion of females that produced a second clutch of eggs and D) number of eggs produced in the post-family life treated females (Ctrl-postFL, N=31 and DM-postFL, N=35). Treatments included ethanol-only exposed control females (Ctrl, grey), and females exposed to Deltamethrin at 6.875 ng/cm² (Delta, orange). Boxplots depict median and interquartile ranges, with whiskers extending to 1.5 times the interquartile range and dots representing jittered experimental values (individual females). ns P > 0.05.

## 4. DISCUSSION

Pesticides are among the most pervasive pollutants in modern agroecosystems, yet their sublethal effects on non-target organisms remain incompletely understood (Desneux et al. 2007; Ndakidemi et al. 2016; Emmerson et al. 2016). While most ecotoxicological studies have focused on survival, reproduction, or overt behavioural changes, these phenotypic endpoints often fail to capture subtle, early, or stage-specific disruptions, as compensatory mechanisms can mask underlying molecular stress. In line with recent calls to complement classical approaches with molecular-level analyses (Lemos et al. 2010; Canzler et al. 2020), we applied a proteomic framework to investigate how sublethal deltamethrin exposure affects protein abundance in field-collected *F. auricularia* females. By sampling individuals at distinct reproductive stages (pre-oviposition and post-family life), we explicitly accounted for contrasting physiological and environmental contexts, including differences in reproductive investment, energy reserves, and metabolic state, which can influence insect responses to pesticide exposure (Bartling et al. 2024). Overall, our results demonstrate that deltamethrin induces both shared and stage-specific proteomic responses, affecting proteins involved in detoxification, metabolism, cytoskeletal organization, and stress responses. Importantly, these molecular changes occurred in the absence of detectable effects on classical reproductive traits, highlighting the capacity of proteomics to reveal cryptic, sublethal impacts that conventional phenotypic assays may overlook.

Our data first show that exposure to deltamethrin produced a consistent proteomic signature across both female reproductive stages, despite the inherent variability of field-collected females in terms of genetic background, developmental history, and physiological state. This shared response involved proteins associated with metabolic regulation and cellular stress. Notably, the upregulation of NADPH–cytochrome P450 reductase, arginine kinase, and glutamate dehydrogenase points to heightened demands on detoxification, energy metabolism, and cellular homeostasis. Conversely, the downregulation of heat shock proteins (HSP70, HSP83) and enzymes involved in lipid metabolism suggests modulation of cellular stress and metabolic processes under chemical exposure. These patterns align with the neurotoxic mode of action of pyrethroids, which induces neuronal hyperexcitation and metabolic imbalance (Clark and Symington 2007; Cao et al. 2011). Similar responses, particularly the regulation of proteins involved in detoxification and energy metabolism, have been reported in several insect species exposed to pesticides (Fujii et al. 2020; Bartling et al. 2024). The consistent regulation of proteins involved in detoxification, metabolism, and cellular stress therefore identifies candidate molecular biomarkers of pesticide exposure in field-collected earwigs. Whether this signature is specific to deltamethrin or reflects a more general response to chemical (or even non-chemical) stressors remains to be investigated. Disentangling this specificity will be critical, as it raises the possibility that this protein network constitutes a central stress-response module, potentially shaping cross-stress outcomes by modulating susceptibility or, conversely, conferring increased resilience to subsequent environmental challenges.

Beyond the shared proteomic signature, our results reveal that deltamethrin exposure also triggered stage-specific responses, highlighting how physiological context modulates the nature and extent of proteomic adjustments to chemical stress. In pre-oviposition females, exposure was associated with the upregulation of proteins involved in fatty acid metabolism, cellular organization and apoptotic processes, while proteins linked to muscle function and calcium homeostasis were downregulated. By contrast, post-family life females exhibited a shift toward upregulation of protein related to protein folding and oxidative stress, alongside a decrease in proteins associated with proteolysis, calcium regulation, and reproductive processes. These differences may reflect the contrasting physiological an environmental condition associated with the two sampling stages, particularly differences in energy reserves and metabolic activity (Zhu et al. 2020; Sharma et al. 2025). Females sampled after the family life stage had experienced prolonged fasting and cold exposure (conditions experimentally implemented to mimic overwintering, a prerequisite for oviposition), which are known to reshape metabolic and stress-response pathways (MacMillan et al. 2016). Within this context, the differential regulation of calcium-related proteins across stages may suggest shifts in neuromuscular or cellular signaling processes, although whole-head extracts preclude distinguishing neuronal from muscular contributions. Notably, several proteins displayed opposite regulation depending on the stage, suggesting that the molecular response to deltamethrin may vary according to the physiological context in which exposure occurs.

In addition to these characterized responses, several uncharacterized proteins were also differentially abundant following deltamethrin exposure. Although their functions remain unknown, their consistent detection across conditions suggests a potential role in the molecular response to chemical stress, possibly in detoxification or associated pathways. These proteins therefore represent promising targets for future investigation, using approaches such as sequence homology analysis, structural prediction, or functional assays (e.g., RNA interference) (Nadzirin and Firdaus-Raih 2012; Xie et al. 2025). Elucidating their roles could uncover previously overlooked mechanisms of pesticide response and adaptation, thereby extending current understanding of how insects cope with chemical stress at the molecular level.

Building on these stage-specific and partly uncharacterized molecular responses, the absence of detectable effects on reproductive traits highlights a key disconnect between molecular disruption and observable phenotype. This finding confirms that proteomic alterations induced by sublethal exposure to pesticides such as deltamethrin do not necessarily translate into measurable changes in life-history traits. One possible explanation is that these molecular perturbations are buffered at the organismal level through compensatory mechanisms that preserve functional stability (Kitano 2004; Campero et al. 2007). Such buffering may conceal underlying physiological costs, which could remain latent and only emerge under additional environmental stressors or over longer timescales, calling for future studies to investigate these long-term effects. Alternatively, the lack of reproductive effects observed here – contrasting with previous studies on the European earwig (Mauduit et al. 2021)-may reflect biological variation among populations or cryptic species within the *F. auricularia* complex, which comprises genetically differentiated lineages with distinct ecological and physiological characteristics (Pasquier et al. 2025a). In this context, differences in population- or species-specific sensitivity to chemical exposure may account for the discrepancies reported across studies, underscoring the importance of considering intraspecific diversity when assessing pesticide effects. Regardless of the underlying mechanism, the proteomic changes detected here indicate that sublethal exposure affects molecular pathways linked to metabolism and reproduction. For instance, the reduced abundance of proteins such as the IGF-II mRNA-binding protein in post-family life females may point to subtle alterations in reproduction, although their functional consequences remain to be determined (Fabrizio et al. 2008).

In conclusion, this study shows that sublethal exposure to deltamethrin is associated with measurable changes in protein abundance in *F. auricularia*, including both shared and stage-dependent responses. These results, obtained from field-collected individuals, indicate that proteomic approaches can detect molecular responses under biologically realistic conditions. The proteins identified here, particularly those related to detoxification and metabolic regulation, may serve as useful candidates for further investigation into biomarkers of pesticide exposure. More broadly, our findings support the idea that molecular perturbations can occur in the absence of detectable phenotypic effects, emphasizing the importance of integrating molecular and organismal approaches to better understand the ecological consequences of chemical exposure.

## ACKNOWLEDGEMENTS

Preprint version 2 of this article has been peer-reviewed and recommended by Peer Community In Ecotoxicology and Environmental Chemistry https://doi.org/10.24072/pci.ecotoxenvchem.100489; Coutellec, M.-A. (2026) Parental care investment modulates insecticide ecotoxicity: an experimental approach in the European earwig.

## FUNDING

This action was led by the Ministries for Agriculture and Food Sovereignty, for an Ecological Transition and Territorial Cohesion, for Health and Prevention, and of Higher Education and Research, with the financial support of the French Office for Biodiversity, as part of “the national call for projects on the Ecophyto II+ plan, part 2, years 2020-2021”, with the fees for diffuse pollution coming from the Ecophyto II+ plan (project *BioIndicFin*). The authors used IA software for English language editing of the manuscript.

## Contributions

Funding acquisition was performed by JM and CL. All authors contributed to the study conceptualization and design. Material preparation and data collection were performed by LP, DT, VL and APTG and statistical analysis were performed by DT and LP. The original draft of the manuscript was written by LP and all authors commented on previous versions of the manuscript. All authors read and approved the final manuscript.

## Ethics declarations

### Ethical approval

Not applicable

### consent to participate

Not applicable

### Consent to publish

All authors approved the final version to be submitted for publication.

### Competing interests

The authors have no relevant financial or non-financial interests to disclose.

## Data availability

The MS proteomics data have been deposited to the ProteomeXchange (Deutsch et al. 2023) Consortium via the PRIDE (Perez-Riverol et al. 2025) partner repository with the dataset identifier PXD076086 and 10.6019/PXD076086. The dataset and the R script have been deposited on Zenodo: https://doi.org/10.5281/zenodo.19912712.

## Notes

https://doi.org/10.5281/zenodo.19912712

### Competing Interest Statement

The authors have declared no competing interest.

### Summary of Updates

Remove the line numbers Remove the page numbers Add the Acknowledgements for recommendation

## REFERENCES

Alexa A, Rahnenführer J (2006) topGO-package: Enrichment analysis for Gene Ontology in topGO. https://bioconductor.org/packages/release/bioc/html/topGO.html. Accessed 17 Sept 2025

Alins G, Lordan J, Rodríguez-Gasol N, et al (2023) Earwig releases provide accumulative biological control of the woolly apple aphid over the years. Insects 14:890. 10.3390/insects14110890

Bartling M-T, Brandt A, Hollert H, Vilcinskas A (2024) Current Insights into Sublethal Effects of Pesticides on Insects. Int J Mol Sci 25:6007. 10.3390/ijms25116007

Beliaeff B, Burgeot T (2002) Integrated biomarker response: A useful tool for ecological risk assessment. Environ Toxicol Chem 21:1316–1322. 10.1002/etc.5620210629

Campero M, Slos S, Ollevier F, Stoks R (2007) Sublethal pesticide concentrations and predation jointly shape life history: behavioral and physiological mechanisms. Ecol Appl Publ Ecol Soc Am 17:2111–2122. 10.1890/07-0442.1

Canzler S, Schor J, Busch W, et al (2020) Prospects and challenges of multi-omics data integration in toxicology. Arch Toxicol 94:371–388. 10.1007/s00204-020-02656-y

Cao Z, Shafer TJ, Murray TF (2011) Mechanisms of pyrethroid insecticide-induced stimulation of calcium influx in neocortical neurons. J Pharmacol Exp Ther 336:197–205. 10.1124/jpet.110.171850

Cheng L, Du Y, Hu J, et al (2015) Proteomic analysis of ubiquitinated proteins from deltamethrin-resistant and susceptible strains of the diamondback moth, *Plutella Xylostella* L. Arch Insect Biochem Physiol 90:70–88. 10.1002/arch.21245

Clark JM, Symington SB (2007) Pyrethroid action on calcium channels: neurotoxicological implications. Invertebr Neurosci IN 7:3–16. 10.1007/s10158-006-0038-7

Desneux N, Decourtye A, Delpuech J-M (2007) The sublethal effects of pesticides on beneficial arthropods. Annu Rev Entomol 52:81–106. 10.1146/annurev.ento.52.110405.091440

Deutsch EW, Bandeira N, Perez-Riverol Y, et al (2023) The ProteomeXchange consortium at 10 years: 2023 update. Nucleic Acids Res 51:D1539–D1548. 10.1093/nar/gkac1040

Dib H, Jamont M, Sauphanor B, Capowiez Y (2011) Predation potency and intraguild interactions between generalist (*Forficula auricularia*) and specialist (*Episyrphus balteatus*) predators of the rosy apple aphid (*Dysaphis plantaginea*). Biol Control 59:90–97. 10.1016/j.biocontrol.2011.07.012

Dietz S, Roman M, Lauck-Birkel S, et al (2009) Ecotoxicological and environmental profile of the insecticide deltamethrin. Bayer Crop J 62:211–226

Diz AP, Martínez-Fernández M, Rolán-Alvarez E (2012) Proteomics in evolutionary ecology: linking the genotype with the phenotype. Mol Ecol 21:1060–1080. 10.1111/j.1365-294X.2011.05426.x

Emmerson M, Morales MB, Oñate JJ, et al (2016) Chapter Two - How agricultural intensification affects biodiversity and ecosystem services. In: Dumbrell AJ, Kordas RL, Woodward G (eds) Advances in Ecological Research. Academic Press, pp 43–97

Fabrizio J, Hickey CA, Stabrawa C, et al (2008) Imp (IGF-II mRNA-Binding Protein) Is Expressed During Spermatogenesis in Drosophila melanogaster. Fly (Austin) 2:47–52. 10.4161/fly.5659

Feng P, Wang Y, Zou H, et al (2023) The effects of glyphosate exposure on gene transcription and immune function of the silkworm, *Bombyx mori*. Arch Insect Biochem Physiol 112:e21990. 10.1002/arch.21990

Fujii T, Sanada-Morimura S, Matsukura K, et al (2020) Energy Reserve Compensating for Trade-Off Between Metabolic Resistance and Life History Traits in the Brown Planthopper (Hemiptera: Delphacidae). J Econ Entomol 113:1963–1971. 10.1093/jee/toaa120

González-Miguéns R, Muñoz-Nozal E, Jiménez-Ruiz Y, et al (2020) Speciation patterns in the *Forficula auricularia* species complex: cryptic and not so cryptic taxa across the western Palaearctic region. Zool J Linn Soc 190:788–823. 10.1093/zoolinnean/zlaa070

Honorio R, Cheutin M –C., Pasquier L, et al (2025) The European earwig: a model species for studying the (early) evolution of social life. Insectes Sociaux 72:177–192. 10.1007/s00040-024-00985-0

Karlsson O, Svanholm S, Eriksson A, et al (2021) Pesticide-induced multigenerational effects on amphibian reproduction and metabolism. Sci Total Environ 775:145771. 10.1016/j.scitotenv.2021.145771

Kitano H (2004) Biological robustness. Nat Rev Genet 5:826–837. 10.1038/nrg1471

Kölliker M (2007) Benefits and costs of earwig (*Forficula auricularia*) family life. Behav Ecol Sociobiol 61:1489–1497. 10.1007/s00265-007-0381-7

Kramer J, Thesing J, Meunier J (2015) Negative association between parental care and sibling cooperation in earwigs: a new perspective on the early evolution of family life? J Evol Biol 28:1299–1308. 10.1111/jeb.12655

Lagadic L, Caquet T, Ramade F (1994) The role of biomarkers in environmental assessment (5). Invertebrate populations and communities. Ecotoxicology 3:193–208. 10.1007/BF00117084

Laufer J, Roche M, Pelhate M, et al (1984) Pyrethroid insecticides: Actions of deltamethrin and related compounds on insect axonal sodium channels. J Insect Physiol 30:341–349. 10.1016/0022-1910(84)90089-1

Le Navenant A, Siegwart M, Maugin S, et al (2019) Metabolic mechanisms and acetylcholinesterase sensitivity involved in tolerance to chlorpyrifos-ethyl in the earwig *Forficula auricularia*. Chemosphere 227:416–424. 10.1016/j.chemosphere.2019.04.065

Lemos MFL, Soares AMVM, Correia AC, Esteves AC (2010) Proteins in ecotoxicology – How, why and why not? PROTEOMICS 10:873–887. 10.1002/pmic.200900470

Lesseur C, Kaur K, Kelly SD, et al (2023) Effects of prenatal pesticide exposure on the fetal brain and placenta transcriptomes in a rodent model. Toxicology 490:153498. 10.1016/j.tox.2023.153498

MacMillan HA, Knee JM, Dennis AB, et al (2016) Cold acclimation wholly reorganizes the *Drosophila melanogaster* transcriptome and metabolome. Sci Rep 6:28999. 10.1038/srep28999

Malagnoux L, Capowiez Y, Rault M (2014) Tissue distribution, characterization and *in vitro* inhibition of B-esterases in the earwig *Forficula auricularia*. Chemosphere 112:456–464. 10.1016/j.chemosphere.2014.05.003

Malagnoux L, Capowiez Y, Rault M (2015a) Impact of insecticide exposure on the predation activity of the European earwig *Forficula auricularia*. Environ Sci Pollut Res 22:14116–14126. 10.1007/s11356-015-4520-9

Malagnoux L, Marliac G, Simon S, et al (2015b) Management strategies in apple orchards influence earwig community. Chemosphere 124:156–162. 10.1016/j.chemosphere.2014.12.024

Martelli F, Zhongyuan Z, Wang J, et al (2020) Low doses of the neonicotinoid insecticide imidacloprid induce ROS triggering neurological and metabolic impairments in *Drosophila*. Proc Natl Acad Sci 117:25840–25850. 10.1073/pnas.2011828117

Mauduit E, Lécureuil C, Meunier J (2021) Sublethal exposure to deltamethrin stimulates reproduction and has limited effects on post-hatching maternal care in the European earwig. Environ Sci Pollut Res 28:39501–39512. 10.1007/s11356-021-13511-7

Merleau L-A, Larrigaldie I, Bousquet O, et al (2022) Exposure to pyriproxyfen (juvenile hormone agonist) does not alter maternal care and reproduction in the European earwig. Environ Sci Pollut Res 29:72729–72746. 10.1007/s11356-022-20970-z

Meunier J, Dufour J, Van Meyel S, et al (2020) Sublethal exposure to deltamethrin impairs maternal egg care in the European earwig *Forficula auricularia*. Chemosphere 258:127383. 10.1016/j.chemosphere.2020.127383

Meunier J, Wong JWY, Gómez Y, et al (2012) One clutch or two clutches? Fitness correlates of coexisting alternative female life-histories in the European earwig. Evol Ecol 26:669–682. 10.1007/s10682-011-9510-x

Müller C (2018) Impacts of sublethal insecticide exposure on insects — Facts and knowledge gaps. Basic Appl Ecol 30:1–10. 10.1016/j.baae.2018.05.001

Nadzirin N, Firdaus-Raih M (2012) Proteins of unknown function in the Protein Data Bank (PDB): an inventory of true uncharacterized proteins and computational tools for their analysis. Int J Mol Sci 13:12761–12772. 10.3390/ijms131012761

Narahashi T, Frey JM, Ginsburg KS, Roy ML (1992) Sodium and GABA-activated channels as the targets of pyrethroids and cyclodienes. Toxicol Lett 64–65:429–436. 10.1016/0378-4274(92)90216-7

Ndakidemi B, Mtei K, Ndakidemi PA (2016) Impacts of synthetic and botanical pesticides on beneficial insects. Agric Sci 07:364. 10.4236/as.2016.76038

Orpet RJ, Crowder DW, Jones VP (2019) Biology and management of European earwig in orchards and vineyards. J Integr Pest Manag 10:21. 10.1093/jipm/pmz019

Pasquier L, Dupont S, Devers S, et al (2025a) Alternative reproductive strategies in two cryptic species of the European earwig complex. Sci Nat 112:48. 10.1007/s00114-025-01999-9

Pasquier L, Groutsch J, Verger M, et al (2025b) Exposure to a glyphosate-based herbicide does not alter maternal care and offspring quality in the European earwig. Ecotoxicology 34:1287–1299. 10.1007/s10646-025-02912-w

Pasquier L, Lécureuil C, Meunier J (2024) Limited effects of a glyphosate-based herbicide on the behaviour and immunity of males from six populations of the European earwig. Environ Sci Pollut Res 31:44205–44217. 10.1007/s11356-024-34063-6

Perez-Riverol Y, Bandla C, Kundu DJ, et al (2025) The PRIDE database at 20 years: 2025 update. Nucleic Acids Res 53:D543–D553. 10.1093/nar/gkae1011

Rehman H, Aziz AT, Saggu S, et al (2014) Systematic review on pyrethroid toxicity with special reference to deltamethrin. J Entomol Zool Stud

Sandrin L, Meunier J, Raveh S, et al (2015) Multiple paternity and mating group size in the European earwig, *Forficula auricularia*. Ecol Entomol 40:159–166. 10.1111/een.12171

Sharma SR, Hwang H, Acharya R, et al (2025) Stage-specific susceptibility of *Bemisia tabaci* MED eggs and neonates to insecticides with different modes of action. Arch Insect Biochem Physiol 119:e70087. 10.1002/arch.70087

Stuligross C, Williams NM (2021) Past insecticide exposure reduces bee reproduction and population growth rate. Proc Natl Acad Sci 118:e2109909118. 10.1073/pnas.2109909118

Van Gestel CAM, Van Brummelen TC (1996) Incorporation of the biomarker concept in ecotoxicology calls for a redefinition of terms. Ecotoxicology 5:217–225. 10.1007/BF00118992

Varela GM, García BA, Stroppa MM (2024) RNA interference of NADPHcytochrome P450 increased deltamethrin susceptibility in a resistant strain of the Chagas disease vector *Triatoma infestans*. Acta Trop 252:107149. 10.1016/j.actatropica.2024.107149

Xie Y, Wang X, Cheng S, et al (2025) RNAi screening of uncharacterized genes identifies promising druggable targets in *Schistosoma japonicum*. PLOS Pathog 21:e1013014. 10.1371/journal.ppat.1013014

Zhang N, Wei J, Jiang H, et al (2022) Knockdown or inhibition of arginine kinases enhances susceptibility of *Tribolium castaneum* to deltamethrin. Pestic Biochem Physiol 183:105080. 10.1016/j.pestbp.2022.105080

Zhu YC, Caren J, Reddy GVP, et al (2020) Effect of age on insecticide susceptibility and enzymatic activities of three detoxification enzymes and one invertase in honey bee workers (*Apis mellifera*). Comp Biochem Physiol Part C Toxicol Pharmacol 238:108844. 10.1016/j.cbpc.2020.108844

