## Supplementary data for "Proteomic signatures of sublethal deltamethrin exposure in the European earwig"

**Table S1. List of the 163 proteins specifically present or absent between Ctrl-preOV and DM-preOV treatments before egg laying.** The accession numbers were obtained from UniprotKB database for Insecta taxonomy.

|  | N° accession | Description |
| --- | --- | --- |
| Present only in  Ctrl-preOV | S5MIA3 | Actin-4 |
|  | A0A288SHR0 | Beta-actin (Fragment) |
|  | A0A084VXA8 | Dper\GL23756-PA-like protein |
|  | A0A6V7HWK5 | Glutamate/phenylalanine/leucine/valine dehydrogenase C-terminal domain-containing protein (Fragment) |
|  | A0AAJ6VP32 | LOW QUALITY PROTEIN: dynamin-like 120 kDa protein, mitochondrial |
|  | A0A484ARM9 | Myosin heavy chain (Fragment) |
|  | A0AA39FEX3 | Ras suppressor protein 1 |
|  | A0A1B1VX07 | Spectrin alpha chain-like protein (Fragment) |
|  | A0A834IDT6 | Small ribosomal subunit protein uS7m |
|  | A0A9J7J109 | Myosin heavy chain, muscle isoform X10 |
|  | A0A6P8ZVJ0 | Myosin heavy chain, non-muscle |
|  | E3UKM1 | Paramyosin (Fragment) |
|  | E0VAJ3 | V-type proton ATPase subunit C |
|  | A0A8I6TI36 | Tubulin beta chain |
|  | A0A8S1D2X0 | Tubulin beta chain |
|  | A0A2J7REM9 | Tubulin alpha chain (Fragment) |
|  | A0A0X9H675 | Tubulin alpha chain (Fragment) |
|  | Q3HLZ8 | Actin (Fragment) |
|  | A0A9N9RZB5 | Actin |
|  | A0A8D8UT74 | Actin, cytoplasmic 1 |
|  | A0A481SZV2 | 60S ribosomal protein L7 |
|  | A0A7R9J7U0 | (California timema) hypothetical protein |
|  | A0A6M2DLR0 | Guanine nucleotide-binding protein subunit alpha |
|  | A0A7R9CBL1 | Small ribosomal subunit protein uS13 |
|  | A0AAD4PFF6 | Polyadenylate-binding protein |
|  | A0A1Y1KCQ5 | 60S ribosomal protein L27 |
|  | A0A1B1VZR3 | Elongation factor 1-alpha (Fragment) |
|  | A0A8K0GF87 | Spectrin beta chain |
|  | A0A9P0CD38 | Moesin/ezrin/radixin homolog 1 |
|  | A0A6J1NWQ2 | Troponin I isoform X11 |
|  | A0A8R1W1S6 | Enoyl-CoA hydratase |
|  | A0A182V429 | Ras-like protein 2 |
|  | A0A6H5HYG8 | Aldehyde dehydrogenase |
|  | A0A6J2Y5D7 | Diacylglycerol kinase |
|  | A0A8D8Q6F9 | Electron transfer flavoprotein-ubiquinone oxidoreductase |
|  | A0A8K0DAD9 | 1-phosphatidylinositol 4,5-bisphosphate phosphodiesterase |
|  | A0A8S4RBU1 | Jg1187 protein |
|  | A0A6V7JX69 | Twitchin (Fragment) |
|  | A0A2L0CAB2 | 60A calcium-transporting ATPase (Fragment) |
|  | A0A2L0CA57 | 60A calcium-transporting ATPase (Fragment) |
|  | A0A6L2PUV4 | Calcium-transporting ATPase |
|  | E2ANL8 | Calcium-transporting ATPase sarcoplasmic/endoplasmic reticulum type |
|  | A0A3Q0J8X5 | P-type Ca(2+) transporter |
|  | K0GEY1 | Sodium/potassium-transporting ATPase subunit alpha |
|  | A0A0A9YG31 | ADP/ATP translocase |
|  | A0A8I6SET0 | Superoxide dismutase [Cu-Zn] |
|  | A0A8S1D587 | SH3 domain-containing protein |
|  | A0A2P1ANK7 | Troponin T |
|  | Q17PR3 | Heat shock protein 83 |
|  | A0A8K0P8Y8 | ATP synthase subunit alpha, mitochondrial |
|  | A0A8K0K0U9 | T-complex protein 1 subunit beta |
|  | A0A1Y1LMP5 | ATP synthase subunit beta (Fragment) |
|  | A0A8I6S4J7 | Catenin delta-2 |
|  | A0A6J1NHV1 | Muscle LIM protein Mlp84B isoform X1 |
|  | A0A336JVI6 | CSON007931 protein |
|  | A0A6P9A9J6 | Protein lap4-like isoform X1 |
|  | A0A835CVP9 | Sterol carrier protein 2 |
|  | A0A7M7Q587 | Prohibitin |
|  | A0A6P9A2H8 | Transforming growth factor beta-1-induced transcript 1 protein |
|  | A0A7R9CN41 | FACT complex subunit |
|  | F5HSC5 | Discs large 1 |
|  | A0A9C6X3S7 | Twitchin isoform X25 |
|  | A0A3B0JGN5 | L-2-hydroxyglutarate dehydrogenase, mitochondrial |
| Present only in  DM-preOV | A0A142I5L4 | Actin (Fragment) |
|  | A0A182N5Q0 | Actin |
|  | A0A9C6TXJ5 | Disco-interacting protein 2 homolog A isoform X1 |
|  | A0A2J7QIQ7 | SH3 domain-containing protein |
|  | A0AAJ7FRH1 | T-complex protein 1 subunit epsilon |
|  | A0AAJ7FP37 | Vesicle-fusing ATPase 1 isoform X1 |
|  | A0A9N9MQT6 | Small ribosomal subunit protein uS2 |
|  | A0A6H5I7C0 | Coatomer subunit beta' |
|  | A0A6P3V2J9 | Protein ERGIC-53 isoform X1 |
|  | A0A1B6DXC2 | Myosin tail domain-containing protein (Fragment) |
|  | A0A1B6KTP5 | Myosin tail domain-containing protein |
|  | T1I795 | Myosin tail domain-containing protein |
|  | A0A0U2P8E2 | Paramyosin |
|  | A0A151WTW5 | Paramyosin |
|  | A0A6I9W5V0 | Paramyosin, long form |
|  | A0A0J7L210 | Succinate--CoA ligase [ADP/GDP-forming] subunit alpha, mitochondrial |
|  | A0A482WUN4 | ATPase V1 complex subunit H C-terminal domain-containing protein |
|  | A0A310SNN2 | Tubulin beta chain |
|  | A0A0L7RCJ9 | Tubulin beta chain |
|  | A0A1W4W2V2 | Tubulin alpha chain |
|  | A0A1S4EA97 | Microtubule-associated protein |
|  | A0A3Q8VIW7 | Actin (Fragment) |
|  | A0A7R9CUY9 | Ribose-phosphate pyrophosphokinase 2 (Fragment) |
|  | A0A7R9H1E6 | CYRIA/CYRIB Rac1 binding domain-containing protein (Fragment) |
|  | A0A1B6HLK6 | RNA helicase |
|  | A0A8K0D737 | RRM domain-containing protein |
|  | A0A7R9FWK0 | PCI domain-containing protein |
|  | A0A2H8TUH5 | Chromatin-remodeling complex ATPase chain Iswi |
|  | A0A1A9US40 | Histone H4 |
|  | A0A8B8HU59 | Ras-related protein Rac1 |
|  | A0A6L2Q579 | Eukaryotic translation initiation factor 3 subunit D |
|  | Q09KA1 | Small ribosomal subunit protein RACK1 |
|  | A0A154PPG8 | DNA ligase |
|  | A0A8S1CA46 | 40S ribosomal protein S3 |
|  | A0A6H5IJ28 | 40S ribosomal protein S3 |
|  | R4FMG4 | Putative ribosomal protein l4 cg5502-pa isoform 1 |
|  | A0A4S2JSX1 | ELAV-like protein 2 |
|  | A0A0N6WGQ4 | Elongation factor 1 alpha (Fragment) |
|  | A0A482X6D0 | Tr-type G domain-containing protein |
|  | A0A1I8QBT0 | Myosin heavy chain, muscle |
|  | W4VRL5 | Putative myosin class i heavy chain |
|  | A0A9R1U7L6 | Dystrophin, isoforms A/C/F/G/H isoform X1 |
|  | A0A0A1XA21 | Tubulin beta chain (Fragment) |
|  | B3P7K9 | GG11120 |
|  | A0A0L7LBA4 | Guanine nucleotide-binding protein alpha-q (Fragment) |
|  | A0A067RL60 | NADH-ubiquinone oxidoreductase 75 kDa subunit, mitochondrial |
|  | A0A8S1CNI1 | UDP-glucose:glycoprotein glucosyltransferase |
|  | A0A6H5J0A4 | Sterol carrier protein 2 |
|  | A0A834KPH2 | Citrate synthase |
|  | A0A7R9HVF7 | Epoxide hydrolase N-terminal domain-containing protein |
|  | A0A9N9SM80 | Serine hydroxymethyltransferase |
|  | A0A9P0TKK9 | 1-phosphatidylinositol 4,5-bisphosphate phosphodiesterase |
|  | A0A0K8TU30 | Malic enzyme (Fragment) |
|  | A0A182V3A8 | Tyrosinase copper-binding domain-containing protein |
|  | A0A9Q0N8Y2 | Growth factor receptor-bound protein 2 |
|  | A0A0A9XVH4 | 2-oxoglutarate dehydrogenase, mitochondrial |
|  | A0A023F5U7 | Putative serine/threonine protein kinase |
|  | A0A9P0XEF4 | pyruvate carboxylase |
|  | A0AAD8DPV1 | arginine--tRNA ligase |
|  | A0A1B0AXE3 | Oxysterol-binding protein |
|  | E0W428 | Rab GDP dissociation inhibitor |
|  | A0A7D9H636 | Clathrin heavy chain (Fragment) |
|  | A0A1L8DVQ7 | Clathrin heavy chain |
|  | A0A2L0CA67 | 60A calcium-transporting ATPase (Fragment) |
|  | A0A2L0C9Y4 | 60A calcium-transporting ATPase (Fragment) |
|  | A0A7R9JU24 | Calcium-transporting ATPase |
|  | K0GG92 | Sodium/potassium-transporting ATPase subunit alpha |
|  | A0A1B6LC86 | ADP/ATP translocase (Fragment) |
|  | A0A1B6L5W3 | ADP/ATP translocase (Fragment) |
|  | A0A8K0G363 | Achaete scute target 1 |
|  | A4GWN4 | Heat shock protein 70 |
|  | A0A5E4Q1D5 | Uncharacterized protein |
|  | W0AMY4 | ATP synthase subunit beta (Fragment) |
|  | A0AA38HGN9 | ATP synthase subunit beta |
|  | A0A8S4FZF7 | (diamondback moth) hypothetical protein |
|  | A0A9C6XSU3 | Protein kinase C and casein kinase substrate in neurons protein 1 isoform X2 |
|  | A0A336MM87 | cysteine desulfurase |
|  | A0AAE1H7K3 | 26S proteasome non-ATPase regulatory subunit 2 |
|  | A0A836G2G4 | MLP2 protein (Fragment) |
|  | A0A7R9JXV1 | Failed axon connections |
|  | A0A6P9A920 | Kinesin light chain |
|  | A0A482WVG3 | NIPSNAP domain-containing protein |
|  | A0A8S4F4B9 | (diamondback moth) hypothetical protein |
|  | A0A1B6IBJ1 | Medium-chain specific acyl-CoA dehydrogenase, mitochondrial (Fragment) |
|  | A0A023F9Y8 | Medium-chain specific acyl-CoA dehydrogenase, mitochondrial |
|  | A0A8B8IHH8 | NADH dehydrogenase [ubiquinone] iron-sulfur protein 3, mitochondrial |
|  | A0A653C6Q9 | Protein transport protein SEC23 |
|  | B2XE08 | Up (Fragment) |
|  | A0A1B6CCK0 | Myosin tail domain-containing protein (Fragment) |
|  | A0A6L2PPX4 | Band 7 domain-containing protein (Fragment) |
|  | G9FUF8 | Alpha-spectrin (Fragment) |
|  | A0A7F5R8F7 | Glycerol-3-phosphate dehydrogenase [NAD(+)] |
|  | A0A0K8TRX0 | Putative sorbin and sh3 domain-containing protein (Fragment) |
|  | A0A6L2P944 | SH3 domain-containing protein |
|  | A0A1J1IJC4 | NADH dehydrogenase [ubiquinone] iron-sulfur protein 3, mitochondrial |
|  | A0A8J6LEK3 | TATA-binding protein interacting (TIP20) domain-containing protein |
|  | A0A482X8A1 | Thioredoxin domain-containing protein |
|  | A0A182W072 | DUF4200 domain-containing protein |
|  | A0AAA9Z4D7 | 2-oxoglutarate dehydrogenase, mitochondrial |
|  | A0A1B6HBE4 | Filamin (Fragment) |

**Table S2. List of the 155 proteins specifically present or absent in DM-postFL compared to Ctrl-postFL treatments after maternal care period.** The accession numbers were obtained from UniprotKB database Insecta.

|  | N° accession | Description |
| --- | --- | --- |
| Present only in  Ctrl-postFL | A0A142I5L4 | Actin (Fragment) |
|  | A0A182N5Q0 | Actin |
|  | A0A7F5RCW6 | Endoplasmic reticulum resident protein 44 |
|  | T1E233 | Putative actin 57b (Fragment) |
|  | A0A2J7QIQ7 | SH3 domain-containing protein |
|  | A0A8W7PMF5 | Uncharacterized protein |
|  | A0A6P3V2J9 | Protein ERGIC-53 isoform X1 |
|  | T1I795 | Myosin tail domain-containing protein |
|  | A0A0U2P8E2 | Paramyosin |
|  | A0A023F9Q6 | Putative succinyl-coa synthetase alpha subunit (Fragment) |
|  | A0AA50XLG9 | Sarco\/endoplasmic reticulum calcium ATPase (Fragment) |
|  | A0A1W4W2V2 | Tubulin alpha chain |
|  | A0A2J7QFW4 | Tubulin alpha chain |
|  | A0A6J0BC13 | Tubulin alpha chain |
|  | A0A3Q8VIW7 | Actin (Fragment) |
|  | A0A1B6DI39 | PDZ and LIM domain protein Zasp |
|  | R4FMG4 | Putative ribosomal protein l4 cg5502-pa isoform 1 |
|  | A0A9N9X8R9 | Insulin-like growth factor 2 mRNA-binding protein 1 |
|  | V5GUB4 | Ubiquitin-ribosomal protein eL40 fusion protein |
|  | A0A6P7GNC1 | Small ribosomal subunit protein uS4 |
|  | A0A482VGF1 | Small ribosomal subunit protein eS25 (Fragment) |
|  | A0A0N6WGQ4 | Elongation factor 1 alpha (Fragment) |
|  | A0A182PQJ0 | Elongation factor 1-alpha |
|  | A0A026WIE2 | Elongation factor Tu |
|  | A0AAD8EDQ5 | Peptidyl-prolyl cis-trans isomerase |
|  | W4VRL5 | Putative myosin class i heavy chain |
|  | A0A1B0CGQ3 | Putative rho guanine nucleotide exchange factor cdep |
|  | A0A9R1U7L6 | Dystrophin, isoforms A/C/F/G/H isoform X1 |
|  | A0A0A1XA21 | Tubulin beta chain (Fragment) |
|  | A0A067RL60 | NADH-ubiquinone oxidoreductase 75 kDa subunit, mitochondrial |
|  | A0A182QUS5 | Aspartate aminotransferase |
|  | A0A1B0FIC6 | Citrate synthase |
|  | A0A9P0TKK9 | 1-phosphatidylinositol 4,5-bisphosphate phosphodiesterase |
|  | A0A154PS73 | Long-chain-fatty-acid--CoA ligase |
|  | A0A0K8TU30 | Malic enzyme (Fragment) |
|  | A0A9Q0N8Y2 | Growth factor receptor-bound protein 2 |
|  | A0A5E4MND2 | Phosphoglycerate kinase |
|  | A0A1L8DVQ7 | Clathrin heavy chain |
|  | A0A2L0C9Y4 | 60A calcium-transporting ATPase (Fragment) |
|  | A0A336KB42 | Calcium-transporting ATPase |
|  | A0A8K0K8A2 | Calcium-transporting ATPase |
|  | K0GG92 | Sodium/potassium-transporting ATPase subunit alpha |
|  | A0AAD9RHA4 | Sodium/potassium-transporting ATPase subunit alpha |
|  | A0A7F5RL76 | ADP/ATP translocase |
|  | A0AAD7ZR24 | T-complex protein 1 subunit theta |
|  | B4N6W0 | ATP synthase subunit beta |
|  | A0A7R8VES6 | Uncharacterized protein |
|  | A0AAE1H7K3 | 26S proteasome non-ATPase regulatory subunit 2 |
|  | A0A836G2G4 | MLP2 protein (Fragment) |
|  | A0A6P9A920 | Kinesin light chain |
|  | A0A1B6IBJ1 | Medium-chain specific acyl-CoA dehydrogenase, mitochondrial (Fragment) |
|  | A0A9P0FK69 | NADH dehydrogenase [ubiquinone] flavoprotein 1, mitochondrial |
|  | A0AAD4K879 | MPN domain-containing protein |
|  | R4G3V1 | Putative 26s proteasome regulatory complex |
|  | A0A0T6BHB1 | Actin-related protein 3 |
|  | A0A653DYN8 | Class II aldolase/adducin N-terminal domain-containing protein |
|  | A9XZN4 | Putative clathrin heavy chain (Fragment) |
|  | A0A7F5R8F7 | Glycerol-3-phosphate dehydrogenase [NAD(+)] |
|  | A0A6L2PP04 | RAVE complex protein Rav1 C-terminal domain-containing protein (Fragment) |
|  | A0A8S0ZN42 | Filamin-A |
|  | N6T3D5 | PX domain-containing protein (Fragment) |
|  | A0AAE1H1I6 | Four and a half LIM domains protein 2 |
| Present only in  DM-postFL | A0A2H4YCD0 | Actin (Fragment) |
|  | T1HZ52 | Actin (Fragment) |
|  | A0A6L2PGW3 | Actin |
|  | A0AAJ7FW76 | Adenosylhomocysteinase |
|  | A0AAJ6VP32 | LOW QUALITY PROTEIN: dynamin-like 120 kDa protein, mitochondrial |
|  | A0AAJ7RST3 | LOW QUALITY PROTEIN: titin |
|  | A0A3Q0IN52 | LOW QUALITY PROTEIN: titin |
|  | A0AAJ6YS57 | LOW QUALITY PROTEIN: twitchin-like |
|  | A0AAJ7N836 | Probable aconitate hydratase, mitochondrial |
|  | A0AA39FEX3 | Ras suppressor protein 1 |
|  | A0A1B1VX07 | Spectrin alpha chain-like protein (Fragment) |
|  | A2I418 | Small ribosomal subunit protein uS19 |
|  | A0A6L2Q299 | Coatomer subunit delta |
|  | A0A1W4XQX5 | Myosin heavy chain, non-muscle isoform X1 |
|  | A0A7R9DAM9 | Myosin motor domain-containing protein |
|  | A0A6M2DP06 | Putative myosin class v heavy chain |
|  | A0A1B6CZ75 | Myosin tail domain-containing protein (Fragment) |
|  | A0A6H5GVT5 | Myosin tail domain-containing protein |
|  | A0A834M359 | Myosin tail domain-containing protein |
|  | A0A0A1XPA0 | Paramyosin, long form |
|  | A0A2P0XJ16 | Putative Per a allergen |
|  | A0A077D5W1 | Sodium-potassium adenosine triposphatase (Fragment) |
|  | E0VAJ3 | V-type proton ATPase subunit C |
|  | A0A8I6TI36 | Tubulin beta chain |
|  | A0A2J7REM9 | Tubulin alpha chain (Fragment) |
|  | A0A8J6HH08 | Na(+)/K(+)-exchanging ATPase |
|  | A0A9J6BYW1 | Dynamin-1-like protein |
|  | A0A0A0WHT2 | Actin (Fragment) |
|  | A0A9N9RZB5 | Actin |
|  | A0A6P4J2W4 | Actin, cytoplasmic type 5-like |
|  | A0A182WJQ3 | Ribosome biogenesis protein WDR12 homolog |
|  | A0A8D8XR91 | rRNA 2'-O-methyltransferase fibrillarin |
|  | A0A9P0QE29 | Histone H2B |
|  | A0A139WF94 | cAMP-dependent protein kinase type II regulatory subunit-like Protein |
|  | A0A1J1IZH1 | Replication factor C subunit 2 |
|  | A0A182IK02 | DNA topoisomerase 2 |
|  | A0A067R0G3 | RNA helicase |
|  | A0A836KDE0 | ADAR editase (Fragment) |
|  | A0AAD4PFF6 | Polyadenylate-binding protein |
|  | A0A7R9AQ25 | U1 small nuclear ribonucleoprotein A |
|  | J3JZK6 | RNA-binding protein lark |
|  | V5GNE3 | 60S ribosomal protein L21 |
|  | A0A182NSP0 | Uncharacterized protein |
|  | A0A5N4AAJ1 | Tubulin beta chain |
|  | A0A8S1CPI4 | Ras-related protein Rab-3 |
|  | A0A2J7R309 | Septin |
|  | A0A034VMJ3 | Bifunctional purine biosynthesis protein ATIC |
|  | T1D424 | Aspartate aminotransferase |
|  | A0A8D8Q6F9 | Electron transfer flavoprotein-ubiquinone oxidoreductase |
|  | A0A6M2DSC2 | Putative gamma-glutamyl kinase (Fragment) |
|  | A0A8S1CRT8 | Glutamate decarboxylase |
|  | A0A9J6BT90 | Glycerol-3-phosphate dehydrogenase |
|  | A0A9P0Q7P5 | Dolichyl-diphosphooligosaccharide--protein glycosyltransferase subunit STT3A |
|  | A0A2J7QWG3 | Pyruvate carboxylase |
|  | A0A3L8DDC9 | Ubiquitin carboxyl-terminal hydrolase 7 |
|  | A0A7R8VG18 | Catenin alpha |
|  | A0A7R9FV54 | Tubulin alpha chain |
|  | E2B894 | Vacuolar protein sorting-associated protein 29 |
|  | A0A2L0CA57 | 60A calcium-transporting ATPase (Fragment) |
|  | A0A482WU54 | Calcium-transporting ATPase |
|  | A0A4D6PJ17 | P-type Ca(2+) transporter (Fragment) |
|  | A0AAE1LMK1 | Vesicle-associated membrane protein 2 |
|  | A0A8I6SET0 | Superoxide dismutase [Cu-Zn] |
|  | A0A8S1D587 | SH3 domain-containing protein |
|  | C7SIR9 | Heat shock protein 70 |
|  | A0A067ZIQ9 | Heat shock protein 70-2 (Fragment) |
|  | A0A6V7GW39 | Heat shock protein 83 |
|  | E0VY10 | Heat-shock protein 105 kDa, putative |
|  | A0A9N9TRI1 | AAA+ ATPase domain-containing protein |
|  | E3W5J8 | Heat shock protein 83 (Fragment) |
|  | A0A2L0ASD9 | Heat shock protein cognate 5 (Fragment) |
|  | A0A7M7Q761 | Heat shock protein 83 |
|  | E0VHJ5 | T-complex protein 1 subunit theta |
|  | A0A4S2KF62 | T-complex protein 1 subunit zeta |
|  | O44097 | ATP synthase subunit beta (Fragment) |
|  | A0A0H5AI05 | ATP synthase subunit alpha, mitochondrial (Fragment) |
|  | W5J415 | Heat shock 70 kDa protein cognate 4 |
|  | A0A8D8JFV1 | Heat shock cognate 71 kDa protein |
|  | A0A8S3WU75 | (apollo) hypothetical protein |
|  | A0A6P3YB46 | Ankyrin-2-like isoform X1 |
|  | A0A6P9A9J6 | Protein lap4-like isoform X1 |
|  | A0A232EZ17 | Coronin |
|  | A0A2J7Q925 | Succinate dehydrogenase [ubiquinone] flavoprotein subunit, mitochondrial |
|  | A0A2J7RDH4 | Paramyosin, short form |
|  | A0A9P0HPU4 | CAP |
|  | A0A7R9HM19 | Uncharacterized protein |
|  | A0A2J7RCF9 | N-terminal methionine N(alpha)-acetyltransferase NatE |
|  | A0A9P0ALB2 | Band 7 domain-containing protein |
|  | A0A3B0JGN5 | L-2-hydroxyglutarate dehydrogenase, mitochondrial |
|  | A0A8K0CG16 | Calponin-homology (CH) domain-containing protein |
|  | A0A194QM76 | Protein Mo25 |
|  | A0A8S9XL47 | Cuticle protein 6 |
|  | A0A1I8PG58 | LIM zinc-binding domain-containing protein |
